# Identification of cereulide producing *Bacillus cereus* by MALDI-TOF MS

**DOI:** 10.1101/324756

**Authors:** Sebastian Ulrich, Christoph Gottschalk, Richard Dietrich, Erwin Märtlbauer, Manfred Gareis

**Affiliations:** Chair for Hygiene and Technology of Milk, Veterinary Faculty, Ludwig-Maximilians-University Munich, Schoenleutnerstr. 8, 85764 Oberschleissheim, Germany; Chair of Food Safety, Veterinary Faculty, Ludwig-Maximilians-University Munich, Schoenleutnerstr. 8, 85764 Oberschleissheim, Germany

**Keywords:** MALDI-TOF MS, *Bacillus cereus*, cereulide, food intoxication

## Abstract

The *Bacillus* (*B*.) *cereus* group is genetically highly homogenous and consists of nine recognized species which are present worldwide. *B. cereus* sensu stricto play an important role in food-borne diseases by producing different toxins. Yet, only a small percentage of *B. cereus* strains are able to produce the heat stable depsipeptide cereulide, the causative agent of emetic food poisonings. To minimize the entry of emetic *B. cereus* into the food chain, food business operators are dependent on efficient and reliable methods enabling the differentiation between emetic and non-emetic strains. Currently, only time-consuming cell bioassays, molecular methods and tandem mass spectrometry are available for this purpose. Thus, the aim of the present study was to establish a fast and reliable method for the differentiation between emetic and non-emetic strains by MALDI-TOF MS. Selected isolates/strains of the *B. cereus* group (total n=110, i.e. emetic n=45, non-emetic n=65) were cultured on sheep blood agar for 48h.

Subsequently, the cultures were directly analyzed by MALDI-TOF MS without prior extraction steps (direct smear method). The samples were measured in linear positive ionization mode in the mass range of *m/z* 800 - 1,800 Da. Using ClinProTools 3.0 statistical software and flex analyst, a differentiation between emetic and non-emetic isolates was possible with a rate of correct identification of 99.1 % by means of the evaluation of two specific biomarkers (*m/z* 1171 and 1187 Da).

**Importance:** *Bacillus* (*B*.) *cereus* plays an important role in food-borne diseases due to the production of different toxins, e.g. the heat stable depsipeptide cereulide. Only a small number of *B. cereus* strains are able to produce this toxin, the causative agent of emetic food poisonings. To minimize the entry of emetic *B. cereus* into the food chain, food business operators require efficient and reliable methods enabling the differentiation between emetic and non-emetic strains. The aim of the present study was to develop a fast and reliable method for the differentiation between emetic and non-emetic strains by MALDI-TOF MS. A differentiation between emetic and non-emetic isolates was possible with a rate of correct identification of 99.1 % by means of the evaluation of two specific biomarkers (*m/z* 1171 and 1187 Da).

## Introduction

*Bacillus* (*B*.) spp. are Gram-positive, rod shaped bacteria occurring world-wide. The *B. cereus* group is genetically highly homogeneous and comprises nine recognized species: *B. anthracis, B. cereus* sensu stricto, *B. cytotoxicus, B. mycoides, B. pseudomycoides, B. thuringiensis, B. toyonensis, B. weihenstephanensis* and *B. wiedmannii* (1, 2). Their spores are heat, acid, UV and desiccation resistant and survive pasteurization (3, 4).

Due to the persistence of spores in food and the ability of the vegetative cells to produce different toxins, isolates of the *B. cereus* group play an important role in food safety. Several publications report an increasing number of cases of foodborne intoxications caused by *B. cereus* in the last years (5, 6). Basically, toxins of the *B. cereus* group can result in two different forms of food intoxication: emetic versus diarrhea (4).

The diarrheal form is caused by heat-labile proteins, i.e. the two enterotoxin complexes Hbl (hemolytic enterotoxin BL) and Nhe (non-hemolytic enterotoxin) as well as cytotoxin K1 (7, 8). In contrast, the emetic form is caused by the small heat-stable cyclic depsipeptide cereulide (Figure 1), encoded by non-ribosomal peptide synthetase genes (*ces*) (9). Cereulide with no structural modification has a molecular mass of 1171 Da (10). Recently, 18 cereulide variants have been described with molecular masses varying from 1147 to 1205 Da. Interestingly, the cytotoxicity of the different analogues differs widely, for instance, the toxicity of isocereulide A is eight times higher than that of cereulide (11).

**Figure 1:**
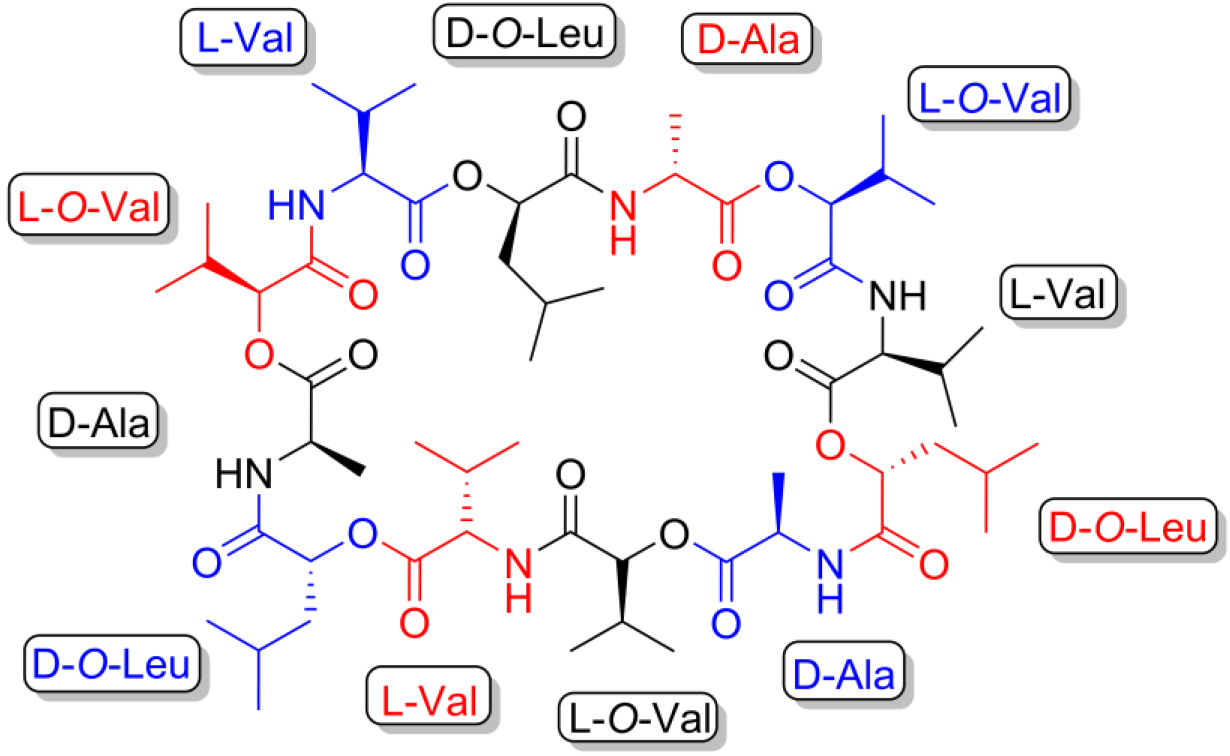
Cereulide, chemical structure of the cyclic dodecadepsipeptide

Emetic *B. cereus* strains represent a major hazard in mass catering and are frequently reported as cause of food-borne outbreaks. Cereulide formation in the digestive tract is rarely observed, if it occurs at all. Intoxication is usually caused by the ingestion of toxin preformed by *ces*-positive *B. cereus* strains in food (12). The direct identification of toxins in food can be carried out by LC-MS/MS or HPLC MS. These methods are able to identify and quantify the toxin with high a sensitivity (10, 13, 14). Generally, food matrices rich in carbohydrates, such as pasta and rice, as well as milk and dairy products have the highest risk of causing cereulide intoxications (6, 15). Symptoms are mainly characterized by vomiting shortly after ingestion of the toxin, with an average duration of one day.

Nevertheless, severe cases require hospitalization and several reports of fatal organ failure have been published (16–20). The detection of viable bacteria prior to toxin production in food is necessary to enable preventive measures in food businesses, such as the elimination of raw materials contaminated with potentially emetic *B. cereus* strains. Therefore, for a general improvement of food hygiene and for the application of specific hazard control plans, a differentiation between emetic and non-emetic *B. cereus* strains would be desirable. However, this requires the availability of a fast and reliable method enabling a high-throughput bacterial screening. Emetic and non-emetic *B. cereus* strains cannot be distinguished by cultural methods used for their isolation (7, 12) and, therefore, mostly molecular methods (21, 22) are used to differentiate *B. cereus* strains based on their genetic profile (9, 23, 24). Furthermore, HEp-2/MTS-bioassays can be used for identifying toxic and non-toxic *B. cereus* strains (25, 26). However, these methods are time consuming, difficult and expensive.

Since some years MALDI-TOF MS is widely used for routine identification of microorganisms (27). This method is extremely robust, fast and suited for operation by laboratory personnel without profound knowledge of the technique per se as the results can be automatically generated via highly sophisticated statistical software. However, previous attempts to apply this technique to identify emetic *B. cereus* were not successful (28). Fiedoruk et al. (28) described a MALDI-TOF MS approach for the differentiation of emetic and non-emetic *B. cereus* strains by measuring proteins in the mass range of *m/z* 4000 - 12000 Da. The MALDI-TOF MS method was regarded as a promising and rapid approach for pre-screening of strains, but was not considered an entirely reliable method to distinguish emetic and non-emetic *B. cereus* strains.

The aim of this study was to establish a MALDI-TOF MS method for a reliable differentiation between emetic and non-emetic *B. cereus* strains by directly measuring the toxin in the biomass obtained by the enrichment of *ces*-positive *B. cereus* cultured on blood-agar.

## Material and methods

### Chemicals

Columbia Agar with 5 % sheep blood was purchased from VWR (Darmstadt, Germany). The MALDI-TOF MS matrix CHCA (*α*-cyano-4-hydroxycinnamic acid), aqua dest. and acetonitrile were purchased from Fluka (Fluka, Dagebüll, Germany). Trifluoracetic acid (TFA) and hydrochloric acid were obtained from Merck (Merck, Hamburg, Germany). Synthetic cereulide standard was purchased from Chiralix (Netherlands), dissolved in ethanol at a concentration of 1 mg/ml and was used for evaluation of the detection limit of the applied MALDI-TOF MS approach.

The matrix for MALDI-TOF MS was prepared according to Meetani and Voorhees (29) by dissolving 10 mg CHCA in 1 ml organic solvent (700 μl acetonitrile, 300 μl aqua dest., 1 μl TFA).

### B. cereus isolates

In total, 110 *B. cereus* strains/isolates were measured and differentiated based on their toxic potential (cereulide production). The isolates were part of the culture collection of the Chair for Hygiene and Technology of Milk (MHI, Department of Veterinary Sciences, LMU Munich). All *B. cereus* were analyzed by molecular methods (PCR assay) and bioassays (HEp-2 cytotoxicity test) for their potential to produce the emetic toxin as described in previous studies (30, 31). In Table 2 the origin of the isolates and their emetic potential are summarized. All isolates were cultured for 48 h at 37 °C on Columbia Agar with 5 % sheep blood before measurement.

### Protein extraction protocol

A small amount (one tip of a wooden application stick) of one colony was transferred directly from the culture medium onto a ground steel target (MTP 384 target plate ground steel BC, Bruker Daltonics GmbH, Bremen, Germany). The spot was overlaid with 1 μl matrix and air-dried at room temperature (approx. 22 °C). After the spot was dried, the sample was again overlaid by 1 μl matrix and allowed to air-dry at room temperature.

### MALDI-TOF MS measurements and data processing

An Autoflex Speed MALDI-TOF/TOF MS (Bruker Daltonics GmbH, Bremen, Germany) was used for measurement. The measurements were performed in linear positive mode (*m/z* 800 - 1,800 Da).

The following parameters were set: random walk of partial sample with ten shots at a raster spot with a limit diameter of 2000 μm; sample rate and digitizer settings were set to 2.00 GS/s, the smartbeam laser was set to “flat” with a frequency of 1000.0 Hz. For automatic measurement an “AutoX” method was created using the “flex control” software. Basic laser settings were laser energy 68 - 78 % (global attenuator offset 24 %) with the following high voltage settings: ion source 1, 19.50 kV; ion source 2, 18.2 kV; lens, 7.0 kV; pulsed ion extraction set to 340 ns. In total, 1000 single spectra per 200 shots were accumulated.

The Bacterial Test Standard (BTS) from Bruker Daltonics GmbH (mass range: 3,637.8 - 16,952.3 Da) was used as calibration standard on a regular basis once a week. With each measurement the peptide standard II from Bruker Daltonics GmbH (mass range: 700 - 3,500 Da) was used for calibration.

All isolates were cultured twice on different days (biological replicates, n=2) and were measured eight times (technical replicates (n=8) per biological replicate). For the differentiation of the emetic and non-emetic strains, the Genetic Algorithm of the statistical analysis software ClinProTools 3.0 (Bruker Daltonics GmbH, Bremen) was applied with the following settings: maximum of peaks: 4; maximum number of generations: 40; mutation rate: 0.2; crossover rate: 0.5. Statistical models were developed with ten emetic (class 1) and ten non-emetic (class 2) strains (n=20, 160 Spectra). For the external validation according to the software, five emetic and five non-emetic strains (n=10, n=80 mass spectra) were processed. Another set of *B. cereus* strains (n=80, 640 Spectra) was applied for classification. For additional quality control all spectra were visually checked for differences in mass peaks with the “flex analyst” software (Bruker Daltoniks GmbH).

To obtain an impression of the sensitivity of the applied MALDI-TOF MS technique a commercially available synthetic standard was analyzed at varying concentrations. For this purpose, the cereulide standard was spotted directly onto the target and, after air drying, was overlaid with the CHCA matrix. The standard was measured in concentrations of 0.001 - 10 μg/ml.

## Results

In a preliminary test the mass spectra of twenty *B. cereus* strains/isolates (ten emetic and non-emetic isolates each) grown on blood agar plates for 48 h were analyzed by MALDI-TOF. The processing of the mass spectra was performed by using statistical models of the ClinProTools 3.0 software. The obtained results, approved the basic applicability of the method for the differentiation of emetic and non-emetic strains (results not shown). To prove the general applicability of the developed method, further 86 *B. cereus* strains/isolates (31 emetic, 55 non-emetic) were analyzed. The analyses were performed as a blind study, i.e. the MALDI-TOF experimenter did not know the assignment of the isolates. Overall, each mass spectrum consisted of approximately 38 discernible mass peaks. Two mass peaks which consistently appeared in all mass spectra of the emetic strains were identified as having prominent intensity differences suitable for the differentiation (*m/z* 1171 and 1187 Da). The emetic strains clearly showed the two mass peaks in their mass spectra, whereas these mass peaks were not detectable within spectra of the non-emetic strains (Figure 2a and b).

**Figure 2:**
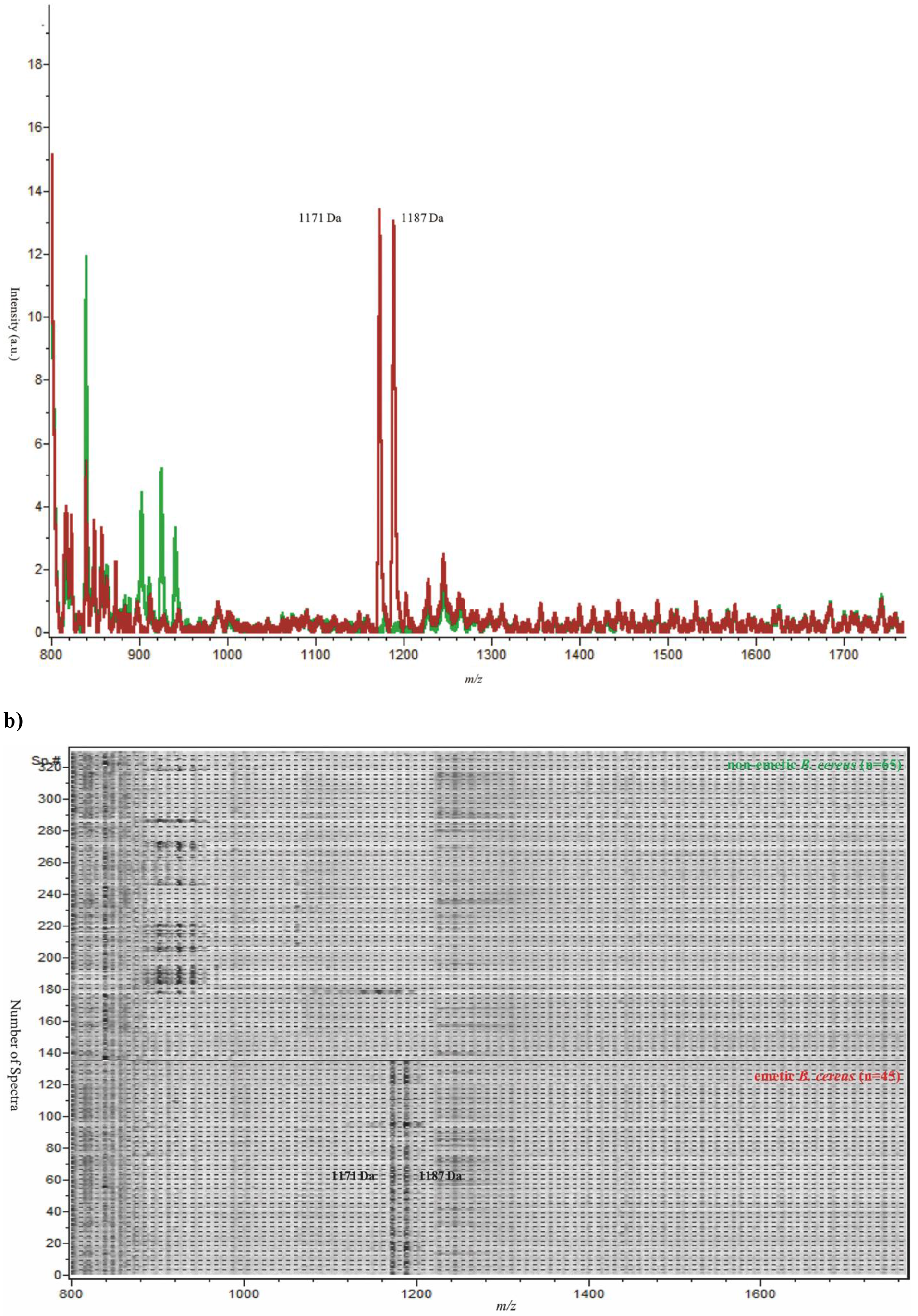
Profile of the 110 analyzed *B. cereus* strains shown as sum spectra (Fig. 2a; emetic strains in red, non-emetic strains in green) or in band view (Fig. 2b). The analyzed *B. cereus* collection comprised 98 isolates, 8 reference strains (see Table 3) and 4 low-producing emetic isolates (see Table 4)

Identification of emetic strains was achieved by evaluation of significant differences (p<0.05) between the average peak intensities of the two different pools (non-emetic class 1 and emetic class 2) which were calculated by t-test/ANOVA and the Receiver Operating Characteristic (ROC values). The mean mass peak intensities ranged from 1.11 - 11.32 a.u. Their ROC values varied from 0.99 – 1.00 AUC (Area Under Curve). Both mass peaks were included into the model showing significant differences in mass peak intensities compared to the non-emetic strains. The calculated difference in average intensities was 5.34 for the *m/z* 1171 Da mass peak and 10.22 for the *m/z* 1187 Da mass peak. The relative standard deviation for the mass peak intensities varied from 0.33 - 6.72 % (Table 1)

**Table 1:**
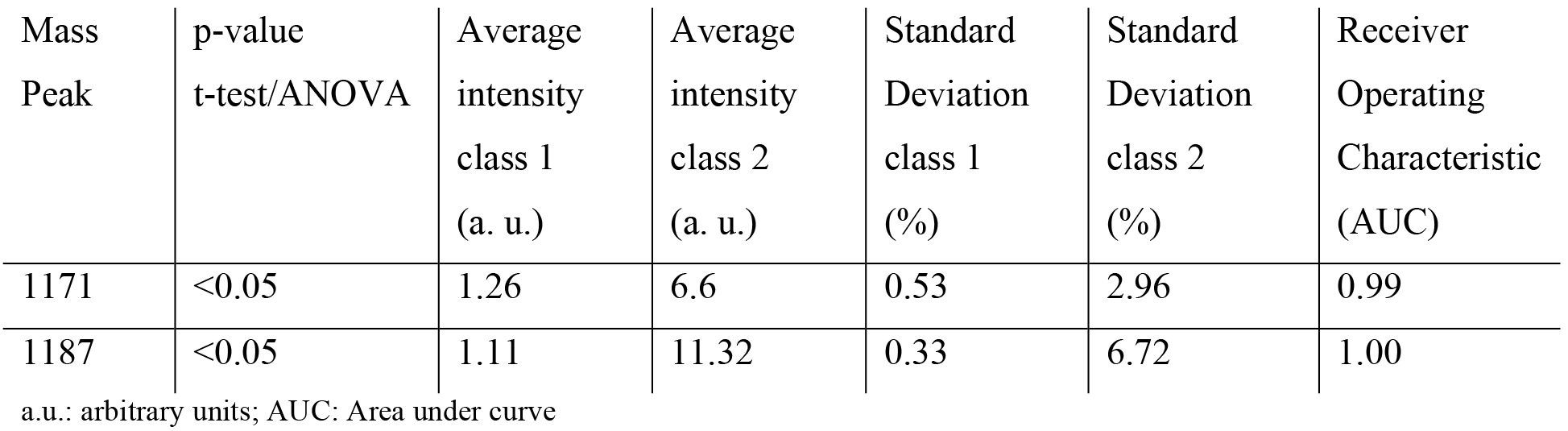
ClinProTools Peak Statistic for the selected mass peaks used for differentiation of non-emetic (class 1) and emetic (class 2) *B. cereus* strains

Overall, the statistical model based on the average mass peak intensity differences of the above mentioned mass peaks enabled the reliable differentiation between emetic (41) and non-emetic (65) strains, including eight reference strains (Table 2 and 3). All mass spectra were also evaluated by visual control for detection of the above-mentioned mass peaks with the software “flex analysis” (Bruker Daltonics GmbH). Altogether, a consistency of 100 % of the MALDI-TOF MS results with the results of prior characterization of the strains was achieved.

**Table 2:**
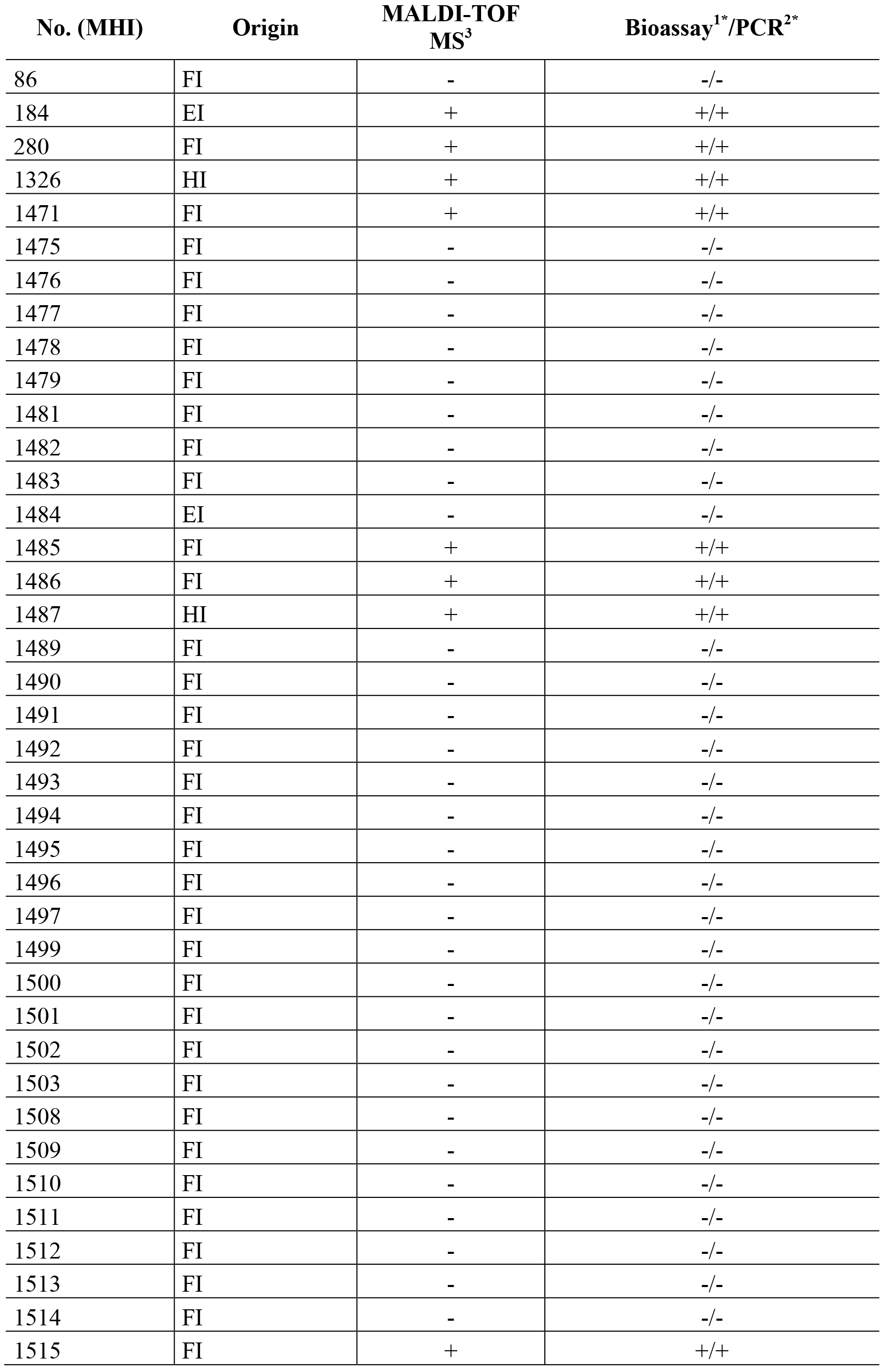
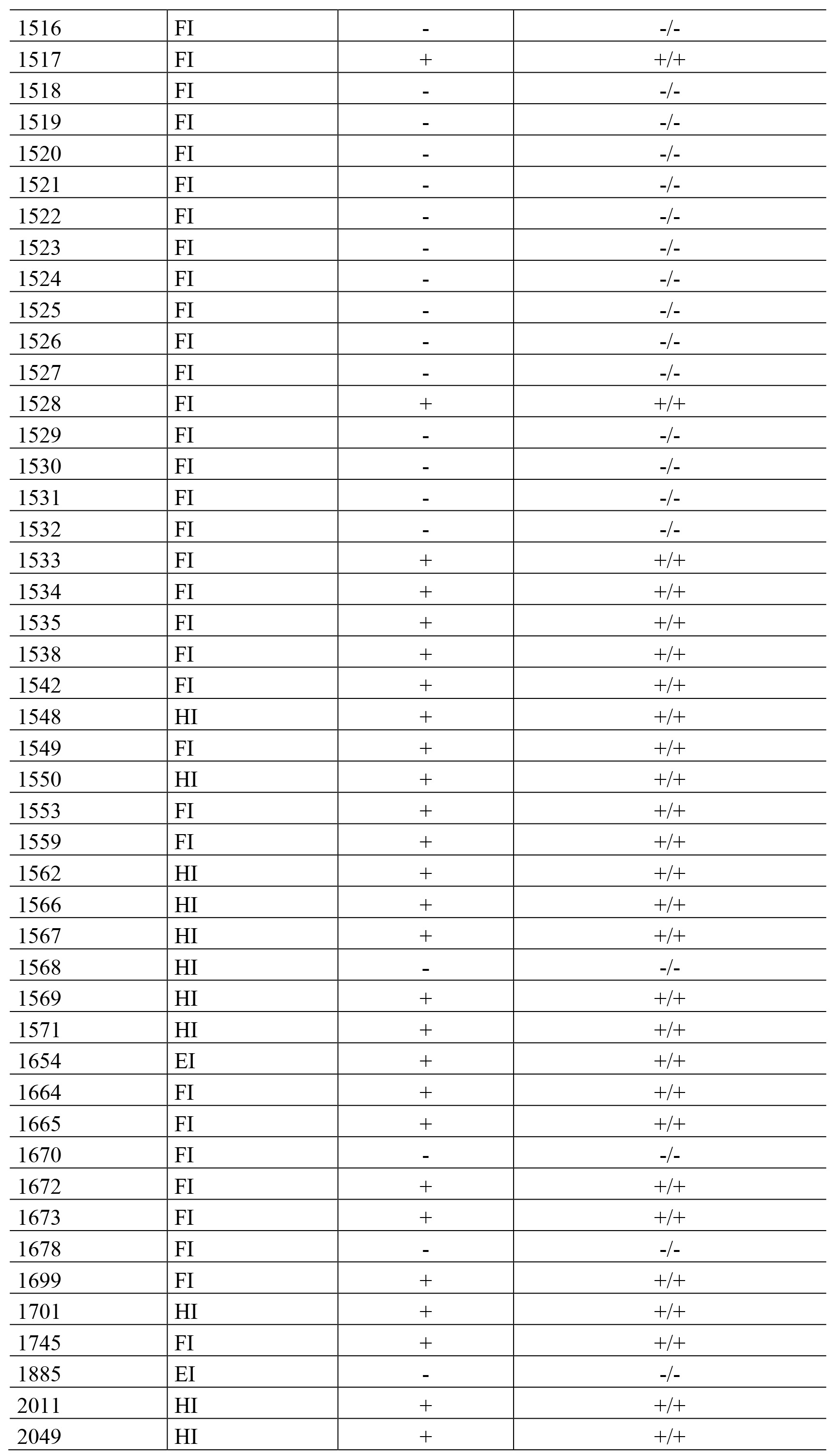
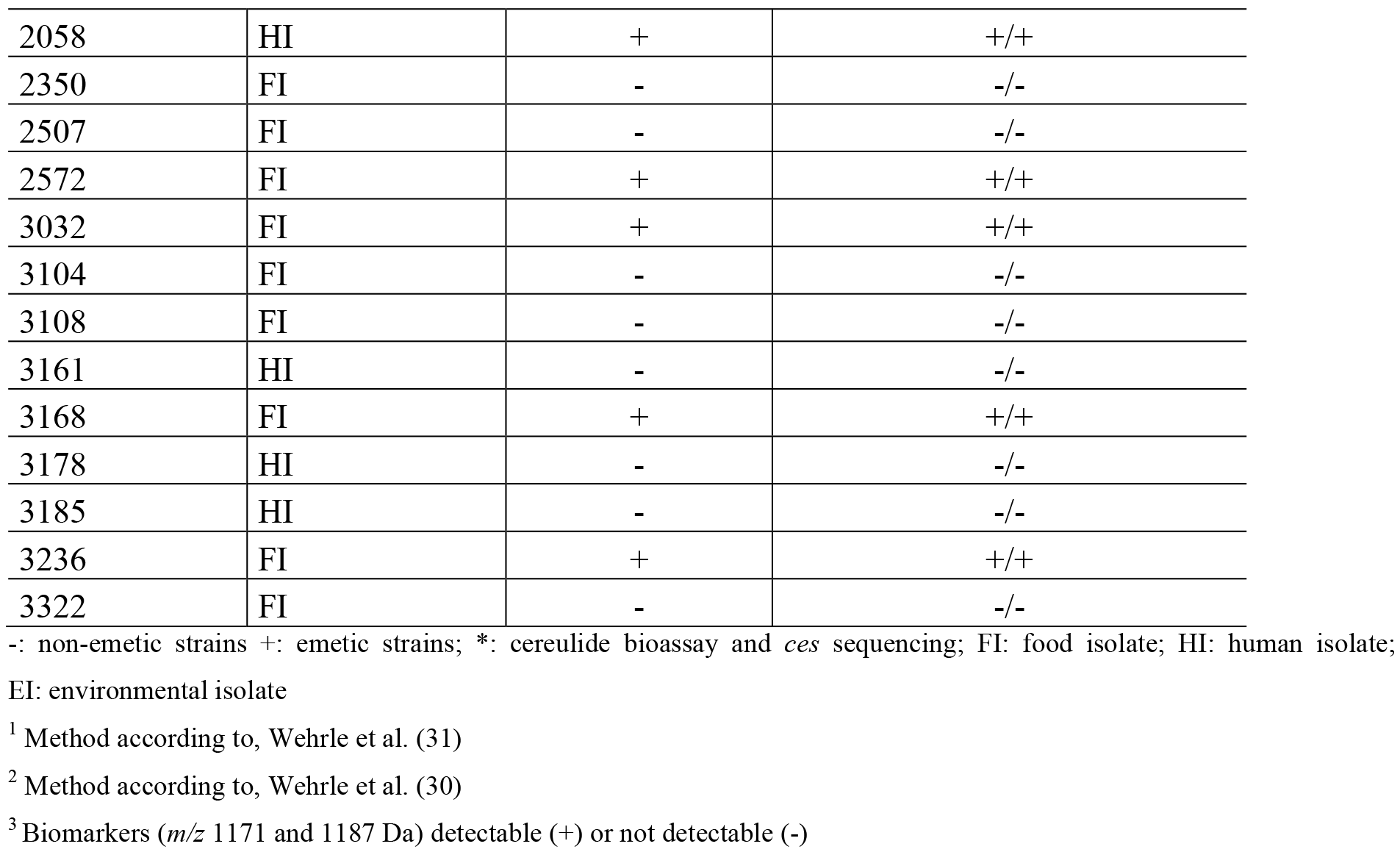
Summary of the differentiation between emetic (+) and non-emetic (−) *B. cereus* isolates by MALDI-TOF MS and comparison with results of previous characterization by bioassays/PCR

**Table 3:**
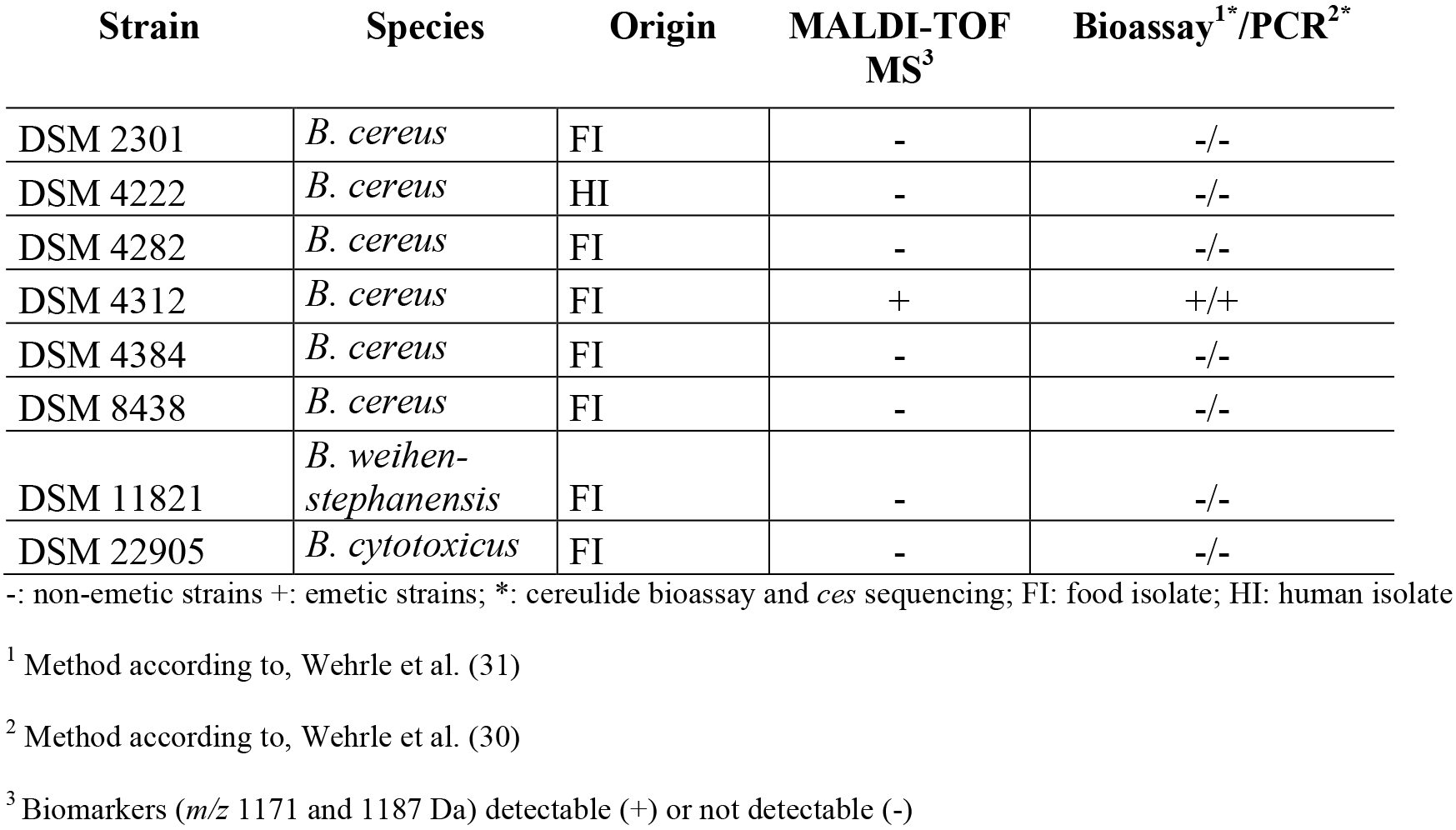
Summary of the differentiation between emetic and non-emetic *B. cereus* reference strains by MALDI-TOF MS and comparison with results of previous characterization by bioassays/PCR

In order to assess the limit of detection for pure toxin, a synthetic cereulide standard was measured in 230 different concentrations (Figure 3). The limit of detection for pure cereulide standard was 1.0 μg/ml when following the described sample preparation protocol.

**Figure 3:**
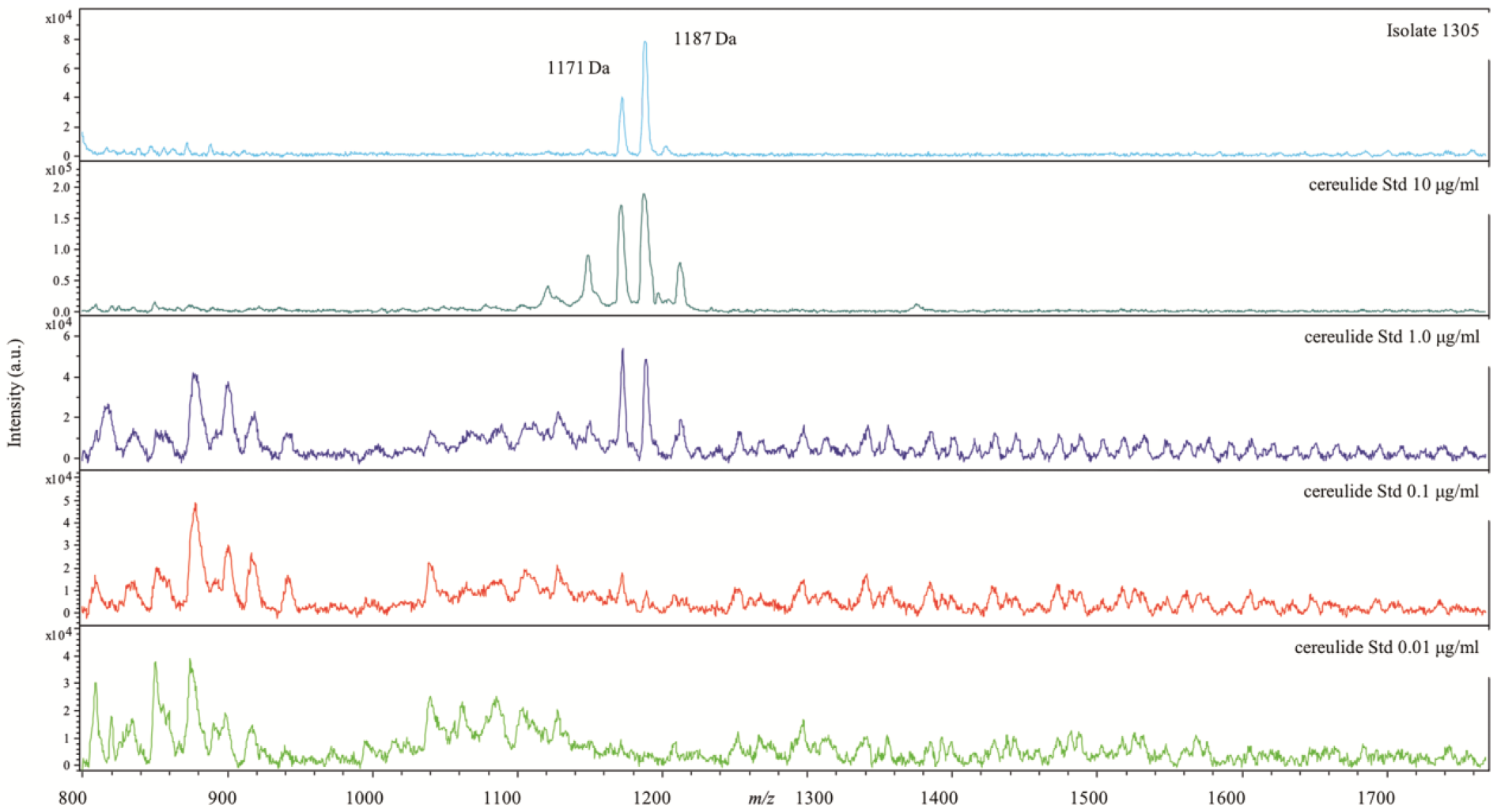
Measurement of synthetic cereulide standard in different concentrations compared to an emetic *B. cereus* strain (MHI 1305)

As several previous studies have pointed out that there is a great variability in the toxin productivity of emetic *B. cereus* strains, we additionally analyzed four low-producing *B. cereus* isolates (Table 4). Apart from the isolate IH 41385 which is well-known for its extremely low productivity, probably caused by a point mutation in the *ces* gene (26), all other analyzed low-producing isolates were correctly identified as emetic *B. cereus* with the established MALDI-TOF technique (Table 4).

**Table 4:**
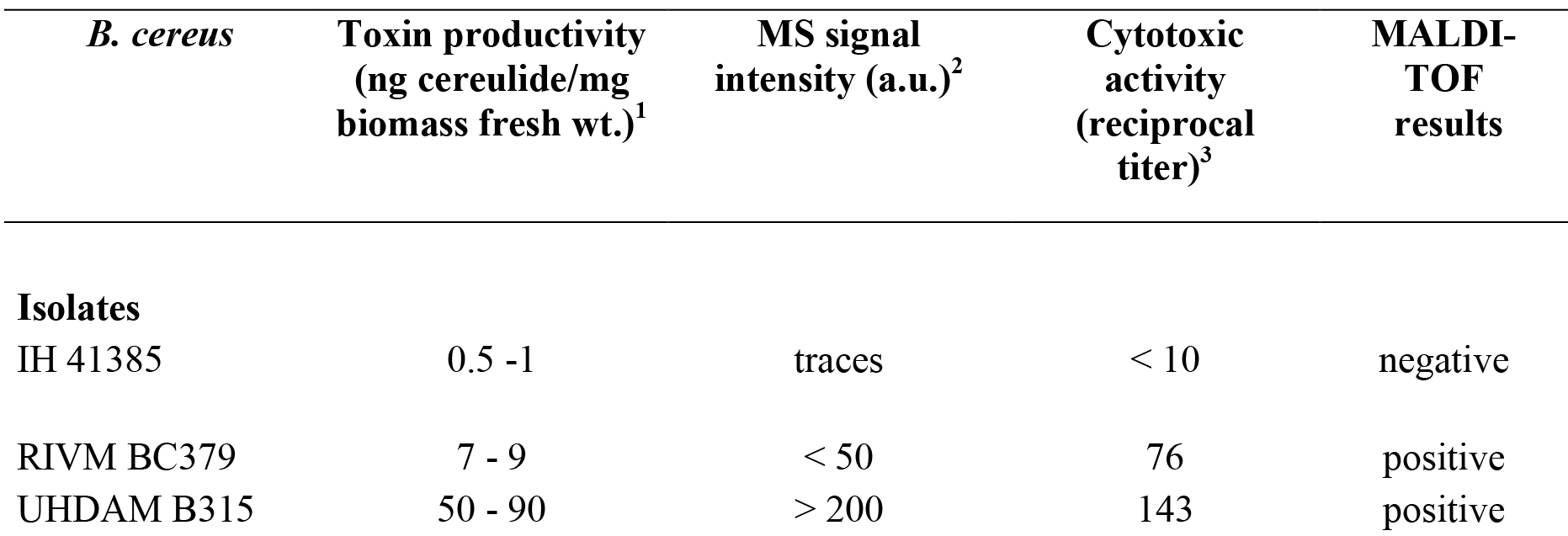
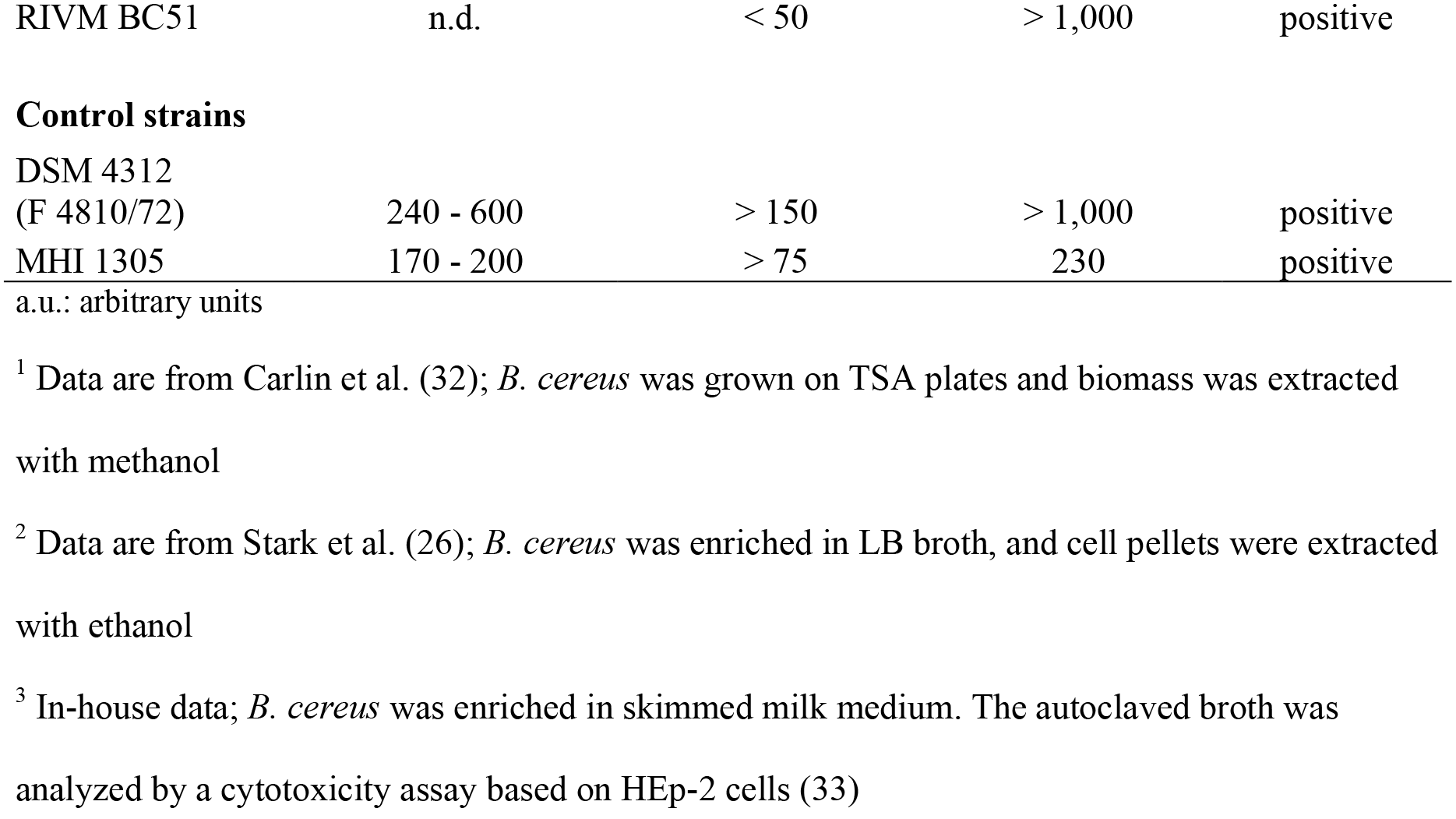
Toxin productivity of low-producing *B. cereus* isolates according to previous publications and in house data. Apart from isolate IH 41385, well-known for its extremely low toxin productivity, all other isolates were correctly identified by MALDI-TOF as cereulide producers.

| *B. cereus* | Toxin productivity<br>(ng cereulide/mg<br>biomass fresh wt.)<sup>1</sup> | MS signal<br>intensity (a.u.)<sup>2</sup> | Cytotoxic<br>activity<br>(reciprocal<br>titer)<sup>3</sup> | MALDI-TOF<br>results |
| --- | --- | --- | --- | --- |
| **Isolates** |  |  |  |  |
| IH 41385 | 0.5 -1 | traces | < 10 | negative |
| RIVM BC379 | 7 - 9 | < 50 | 76 | positive |
| UHDAM B315 | 50 - 90 | > 200 | 143 | positive |
| RIVM BC51 | n.d. | < 50 | > 1,000 | positive |
| Control strains |  |  |  |  |
| DSM 4312<br>(F 4810/72) | 240 - 600 | > 150 | > 1,000 | positive |
| MHI 1305 | 170 - 200 | > 75 | 230 | positive |

a.u.: arbitrary units
<sup>1</sup> Data are from Carlin et al. (32); *B. cereus* was grown on TSA plates and biomass was extracted with methanol
<sup>2</sup> Data are from Stark et al. (26); *B. cereus* was enriched in LB broth, and cell pellets were extracted with ethanol
<sup>3</sup> In-house data; *B. cereus* was enriched in skimmed milk medium. The autoclaved broth was analyzed by a cytotoxicity assay based on HEp-2 cells (33)

## Discussion

*B. cereus* is abundant in the environment and, thus, is frequently found in food. Low levels of *B. cereus* cells or spores are found on virtually every raw agricultural commodity. Particularly, herbs and other food material that have direct contact with soil are at high risk for contamination with *B. cereus* (7, 34). Due to their food poisoning potential higher levels of *B. cereus* in food constitute a public health hazard and represent a major problem for the food industry. This applies particularly to the emetic strains capable of producing high amounts of the heat-stable cereulide in food. Depending on the food category investigated, prevalence rates for emetic strains show a broad variability, percentages in the range from <1 % in vegetables to >20 % in farinaceous products have been reported (12, 15, 35). To improve HACCP based concepts and prevent foodborne intoxications by emetic *B. cereus* it is necessary to identify the currently unknown entrance-points into the food production (6). This requires novel diagnostic strategies as morphological or microscopic approaches are nearly useless for the differentiation of emetic and non-emetic *B. cereus* strains (33). Identification of emetic strains is currently only possible by complex and sophisticated methods such as PCR, bioassays or mass spectrometry (6, 14, 23, 26, 28).

In contrary, MALDI-TOF MS can be used as a fast screening method for routine microbiological analysis since minimal sample pretreatment is required. Therefore, this technique appears quite as a preferable method for the differentiation of emetic and non-emetic strains. However, up to now, only one study has been published in which the applicability of this technique to this purpose was evaluated (28). In principle the authors used an indirect approach by measuring differences in the mass spectra profiles in positive linear ionization mode in the range of *m/z* 4000 – 12000 Da. Ultimately, after analyzing more than 100 *B. cereus* isolates, the authors stated that proteomic profiling of whole cells by MALDI-TOF MS is not a sufficiently reliable method to distinguish emetic and non-emetic *B. cereus*.

Therefore, in the present study a direct approach to differentiate between emetic and non-emetic strains was chosen. Obviously, the direct approach has the limitation of being dependent on the production of cereulide on the culture medium and the temperature used for cultivation. If a strain has the ability to produce cereulide but does not produce cereulide on an agar plate, the result of the MALDI-TOF MS measurement would be false negative. In principle, like previously described for many other bacterial toxins, cereulide production depends strongly on the growth and enrichment conditions applied (12, 36). For the evaluation of the toxin productivity of emetic *B. cereus* strains, in earlier studies the bacteria were grown on tryptic soy agar (TSA) plates for up to ten days at 28 °C and then the biomass was extracted by organic solvents (37). A more rapid approach was used by Stark et al. (26) in which isolates were precultured in LB broth (TSB) and then enriched overnight at 24 °C. However, the subsequent extraction of the cell pellet took up to 17 h. While both these culture media resulted in high toxin amounts, no toxin productivity could be observed in other media such as BHI and peptone broth commonly used for the enrichment of bacteria (38). Less known is that blood agar plates also represent an excellent medium for cereulide production (39). Therefore, in our approach, bacterial cultures were directly measured from the blood agar plate and the detection of cereulide positive/negative samples could be performed within 5 minutes.

Comparing the detected mass peaks of the cereulide standard (Figure 3) with the mass peaks (*m/z* 1171 and 1187 Da, Figure 2) found after analyzing whole cells of emetic strains, it is fairly certain that these mass peaks represent the cereulide produced by the strains. Overall, the method worked very well, 109 out of 110 tested *B. cereus* strains/isolates were correctly identified. Only one emetic strain (IH 41385, Table 4) reacted false negative in the MALDI-TOF MS. This particular strain is well-known for its extremely low productivity (26, 32). Considering the toxin dosis needed to induce emesis, i.e. 8 μg cereulide per kg body weight (40), it is highly unlikely that such low-producer strains represent a public health hazard.

In conclusion, the developed method is characterized by a high inclusivity of >99 % and a striking simplicity. Theoretically a strain may produce cereulide in a food matrix but not on a blood agar plate (41). However, in our analyses including a comprehensive range of emetic *B. cereus* strains from different sources we found no indication for this scenario. Whether the method is equally suited for detection of cereulide in other bacteria of the *B. cereus* group, e.g. *B. weihenstephanensis,* has to be further investigated (42, 43). Future research may also reveal if a modified version of the presented method is additionally applicable for the identification of the diarrheal toxins, i.e. Hbl or Nhe produced by *B. cereus.*

## Acknowledgements

Sponsored by the Adalbert Raps foundation, Kulmbach, Germany. Any opinions expressed here are those of the authors.

